# High precision coding in visual cortex

**DOI:** 10.1101/679324

**Authors:** Carsen Stringer, Michalis Michaelos, Marius Pachitariu

**Affiliations:** HHMI Janelia Research Campus, Ashburn, VA, USA

## Abstract

Single neurons in visual cortex provide unreliable measurements of visual features due to their high trial-to-trial variability. It is not known if this “noise” extends its effects over large neural populations to impair the global encoding of stimuli. We recorded simultaneously from ∼20,000 neurons in mouse primary visual cortex (V1) and found that the neural populations had discrimination thresholds of ∼0.34° in an orientation decoding task. These thresholds were nearly 100 times smaller than those reported behaviorally in mice. The discrepancy between neural and behavioral discrimination could not be explained by the types of stimuli we used, by behavioral states or by the sequential nature of perceptual learning tasks. Furthermore, higher-order visual areas lateral to V1 could be decoded equally well. These results imply that the limits of sensory perception in mice are not set by neural noise in sensory cortex, but by the limitations of downstream decoders.

---

Sensory neurons respond with high variability to repeated presentations of the same stimulus ^1;2;3;4;5;6;7;8^. This variability is thought to limit the accuracy of perceptual judgements because it is pervasive ^9;10;11;12^, and it is reduced during attentional engagement ^13;14;15^ and over the course of perceptual learning ^16^. The hypothetical links between neural variability, sensory information and perception form the foundations of several theoretical frameworks such as the efficient coding hypothesis ^17;18^, the information bottleneck ^19^, the ideal observer model ^20^, the Bayesian coding hypothesis ^21;22;23^ and the probabilistic sampling hypothesis 24;25.

However, it is not clear how variability measured from single neurons or from pairs of neurons scales to local circuits of tens of thousands of neurons ^10^. Intuitively, one might expect the noise to be averaged out. Theoretical studies show that most types of noise are indeed harmless at the population level ^26;27;28;29^; only a special kind of correlated noise is detrimental to neural coding because it limits the total information available in the system. This “information-limiting” noise arises when the estimation errors of single neurons are correlated to each other across the population ^29^.

In geometric terms, noise only affects the encoding of a stimulus when it aligns to the same neural sub-spaces which the stimuli drive ^30;31;32^. Previous studies have shown that at least some of the neural variability, that induced by the animal’s own behavior ^33^, is orthogonal to the stimulus subspace and thus harmless. However, such behavior-related variability accounts for only ∼35% of the population-level variability, leaving the possibility that the rest is stimulus-related and thus potentially information-limiting.

Estimating the impact of information-limiting noise on coding is difficult, because even small amounts of such noise can put absolute bounds on the precision of stimulus encoding ^29^. To detect the effects of potentially small but information-limiting noise, it is thus necessary to record from large numbers of neurons during a large number of trials. To estimate the information content of such multi-neuron recordings, we used decoding approaches. Previous studies in anesthetized macaques show that one-dimensional variables, like the orientation of a grating, can be decoded from populations of 10-100 simultaneously-recorded neurons with errors in the 3-20° range ^34;35;36^. It is not known if these errors represent an absolute lower bound, or may decrease further for larger neural populations. Some studies claim that small subsets of neurons are as discriminative as the entire population ^35;37^, while others show consistent improvements in decoding with increasing numbers of neurons ^38^. It is also not known if having more training trials would reduce overfitting and thus allow the decoding of even smaller orientation differences; previous studies were limited to stimulus densities of 4-10 trials per degree ^35;36^.

Here we aimed to determine if neural noise puts fundamental limits on stimulus encoding accuracy, by recording from populations of ∼20,000 neurons in mice, using stimulus sets with densities of up to 1,000 trials/degree. If information-limiting noise exists, the decoding error should asymptote at some non-zero value as we increase the number of neurons and trials we consider ^29^. We found that the error did not asymptote, and the discrimination thresholds were as low as 0.3°. To achieve this decoding performance, we had to use decoders that take correlations into account. Thus, the visual cortex encodes stimuli to high precision on a trial-by-trial basis in mice, a species not known for high acuity vision, and which performs poorly in orientation discrimination tasks ^39;40;41^.

## Results

We recorded from primary visual cortex in awake, head-fixed mice that were free to run on an air floating ball. Each session lasted for 120-180 minutes during which we presented images to the left eye (Figure 1a). Our main stimuli were static gratings rotated at a random orientation on each trial, which lasted for 750ms, followed by 500 ms of gray screen. We recorded neural activity from visual cortex using multi-plane two-photon calcium imaging, with 10-17 planes spaced 25 *µ*m apart in depth, scanning the entire stack repeatedly at an average 2.5-4 Hz (Figure 1b). For the main stimulus set, we obtained 19,665 ± 3,062 (s.d., n=6 animals) neurons per recording using the processing pipeline Suite2p ^42^ (Figure 1c and Movie 1). All analyses were performed on deconvolved data, which localizes in time the fluorescence responses of the calcium indicator ^43^. The responses to each stimulus were defined as the average, deconvolved fluorescence over the 3 bins following stimulus onset. We have publicly shared the data and code for this paper ^44^.

**Figure 1:**
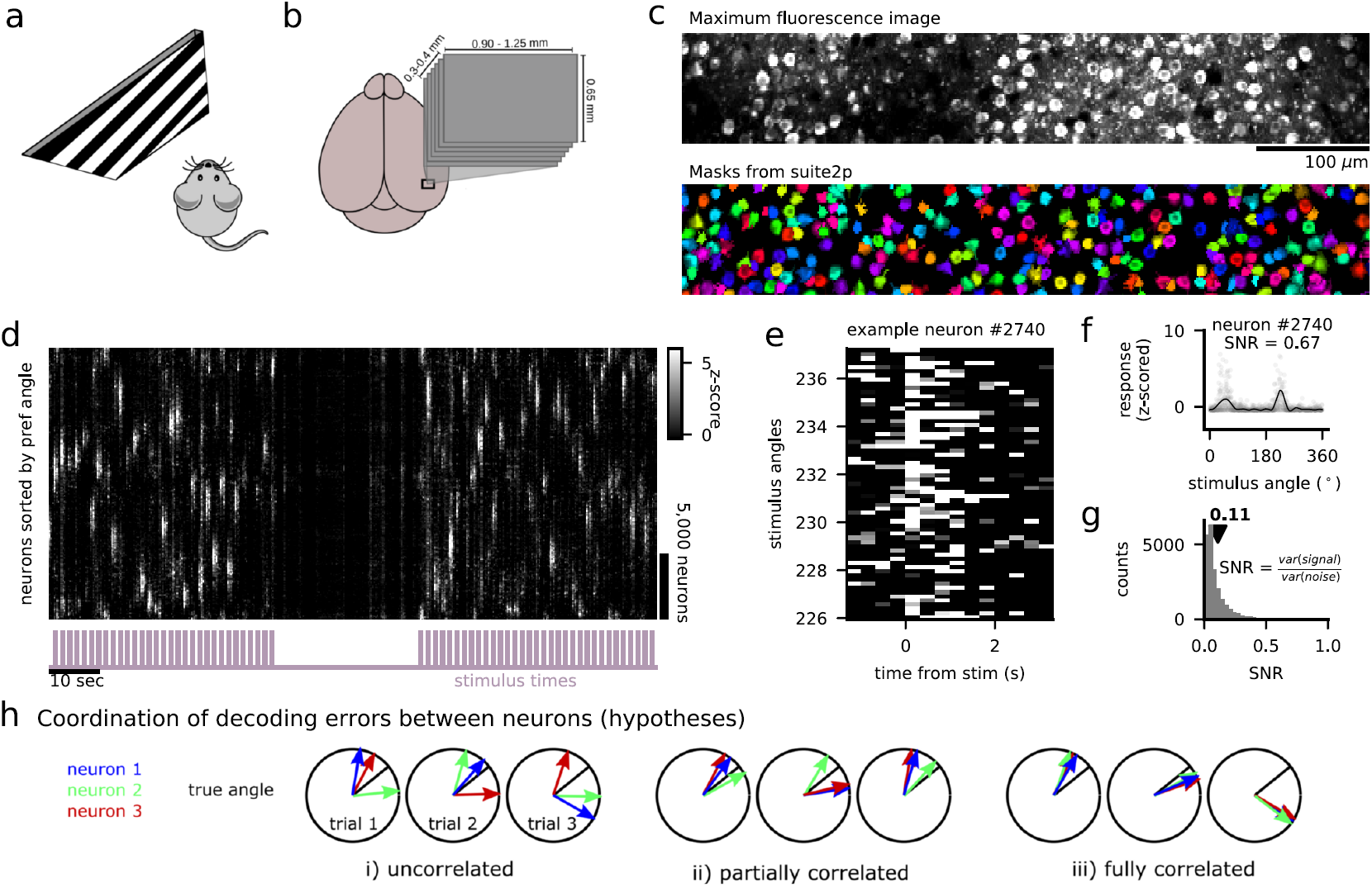
Recording setup and single-neuron variability. **a**, Visual presentation setup. **b**, Multi-plane imaging setup. **c**, Section from a recorded plane. (top plot) maximum fluorescence image. (bottom plot) cell masks detected with Suite2p, randomly colored. **d**, Activity raster of all neurons in one recording in response to static gratings of various angles. Neurons are sorted along the vertical axis by their preferred angle, and smoothed with a Gaussian of width 50 neurons. **e**, Single trial stimulus responses of an example neuron. **f**, Responses from **e** averaged over 3 bins following stimulus onset (gray dots), with average tuning curve (black line). **g**, Distribution of the signal-to-noise ratio (SNR) across neurons in this session. **h**, Hypotheses for the alignment of decoding errors across the neural population. Each circle represents a different trial. The black line denotes the true angle of the stimulus. Each arrow represents the decoded angle from a single neuron’s response. (i) the errors are uncorrelated across neurons. (ii) some errors are correlated. (iii) all errors are correlated, and therefore information about the stimulus is lost.

To visualize the patterns of population activity in a raster plot, we sorted neurons by their preferred stimulus (Figure 1d). As previously shown ^45^, single neurons had high trial-to-trial variability and sometimes failed to respond at all to their preferred stimulus (Figure 1e and Movie 1). We quantified this variability by the signal-to-noise ratio (SNR) (Figure 1f), and found a mean SNR of 0.11 for the example session, and 0.13 ± 0.01 (s.e.m., n=6) across recordings, similar to the single-trial SNR previously reported for natural image stimuli ^38^. We also compared the single-neuron, single-trial signal variance from our two-photon datasets to that from electrophysiological recordings (0.11 vs 0.12) and found that there was little loss of information due to our recording method (Figure S1a,b) ^46^. The aligned, population-averaged tuning curves had a mean half-width at half-max of 14.1°(Figure S1d) ^47^. The correlation between neural response vectors for different orientations decayed smoothly as a function of the difference in orientation (Figure S1g,h). Using a manifold embedding algorithm in three dimensions (ISOMAP ^48^), we found a predominantly one-dimensional representation of the stimulus space, but nonetheless a representation corrupted with noise (Figure S1i).

We distinguish between three possible types of neural variability that affect coding in different ways (Figure 1h). First, the decoding errors of single neurons might be uncorrelated, in which case averaging over single-neuron predictions would give an unbiased estimate of the true stimulus angle with a small error (Figure 1h(i)). A second possibility is that decoding errors are partially-correlated, for example if subsets of neurons have correlated errors, in which case averaging would not be as effective and the optimal decoder needs to take correlations into account (Figure 1h(ii)). The third and final possibility is that decoding errors are fully correlated over the population, so that averaging over the predictions of even infinitely many neurons would give a biased estimate of the true stimulus orientation (Figure 1h(iii)). This situation would indicate the presence of information-limiting correlations ^29^.

### Independent neuron decoder

Consider the first possibility, that the stimulus-dependent variability is uncorrelated between neurons. If that was true in the data, we could decode from the neural population using the “Naive Bayes” classifier ^49^, which applies Bayes’ theorem under the assumption of independence between neurons. Given a response *R*(*n,t*) of neuron *n* on trial *t*, we estimated the probability P(θ|*R*(*n,t*)) ∼ *𝒩* (*R*(*n, t*)|*f*_*n*_(θ), σ_*n*_), for every possible angle θ (Figure 2a). On training data, we derived *f*_*n*_(θ) and σ_*n*_ as the mean tuning curve and the variance of the neural response, respectively (Figure 2a). The Naive Bayes classifier multiplies these probabilities across neurons, or equivalently *sums* the *log*-likelihoods (Figure 2a). Finally, the stimulus with the highest summed probability is selected as the decoded orientation, using interpolation to predict fractional stimulus values (see Methods).

**Figure 2:**
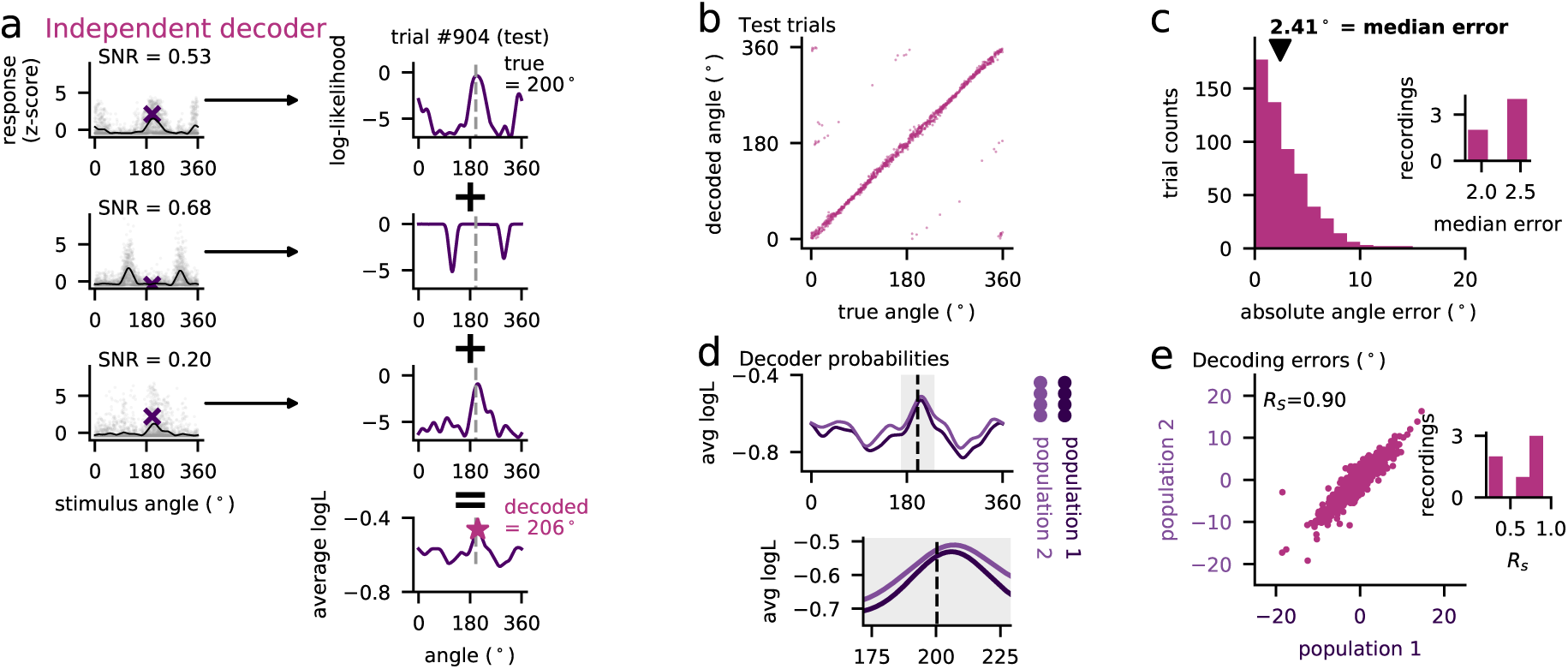
Independent decoders of stimulus orientation. **a**, Single-neuron single-trial stimulus responses (gray dots) with average tuning curves (black line). Right panel: log-likelihood of stimulus orientation based on each neuron’s observed response on trial #904. Actual stimulus is gray dotted line. Right, bottom panel: average log-likelihood over all neurons. **b**, True versus decoded stimulus on test trials. **c**, Distribution of absolute angle errors across trials. (inset: distribution of median decoding errors across recordings). **d**, Neurons were split into two random populations and the decoder was fit separately to each population. **e**, Scatter plot of decoding errors from the two populations. The Spearman correlation *RS* is reported (inset: distribution of *RS* across recordings).

This population decoder had a median error of 2.31 ± 0.14°(s.e.m., n=6 animals) (Figure 2b,c). This error may be either due to single-neuron noise that was not fully averaged out, or due to correlations in decoding errors between neurons. To distinguish between these two scenarios, we split the neurons into two populations, decoded from each, and asked if their decoding errors were in the same direction with respect to the true stimulus (Figure 2d). We found that the errors were highly correlated (Spearman’s R = 0.65 ± 0.12, s.e.m, n=6), which invalidated the independence assumption of the Naive Bayes decoder (Figure 2e). This suggests that the error may decrease further if correlations are taken into account, which we show in the next section.

### Linear decoders account for correlations

To account for correlations, a decoder must be able to appropriately weigh neurons, potentially discarding neurons that have correlations which are detrimental to coding. This is easily achieved with simple linear decoders, which we trained to predict continuous functions of the stimulus orientation (Figure 3a). We call these intermediate functions “super-neurons” and chose them to have von Mises tuning to orientation θ (Figure 3a): *F*(θ|θ_pref_) = exp(cos(θ − θ_pref_)*/*σ) with σ = 0.1. Different super-neurons had different values of θ_pref_ that tiled the full range [0, 2π]. On test data, the decoded stimulus was the preferred angle of the super-neuron with the highest activation, using interpolation methods to decode fractional angles (Figure 3a).

**Figure 3:**
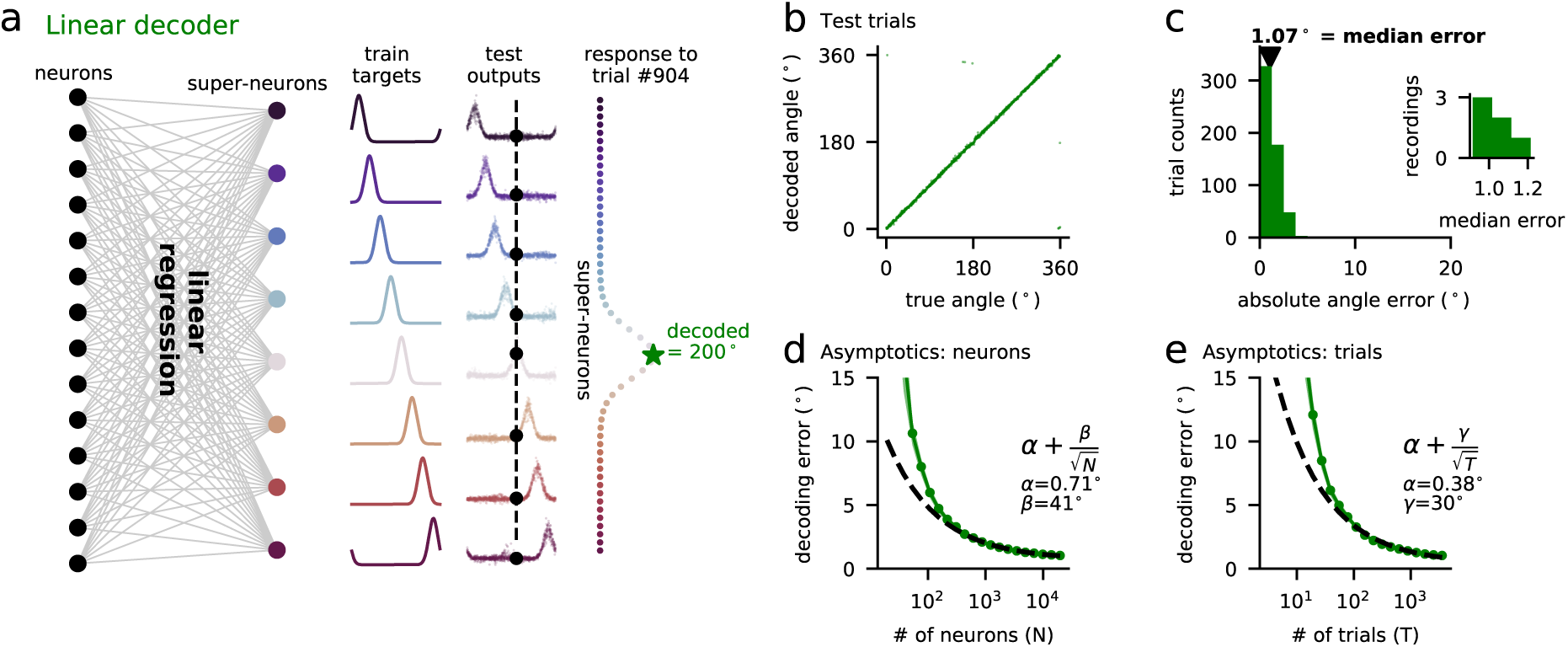
Linear decoders of stimulus orientation. **a**, Schematic of linear decoder, with “super-neuron” responses to test stimuli (same example trial as Figure 2a). **b**, True versus decoded stimulus on test trials. **c**, Distribution of absolute angle errors across trials. (inset: distribution of median decoding errors across recordings). **d**, Scaling of decoding error as function of the number of neurons considered, together with fitted curve 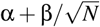 where *N* is the number of neurons. **e**, Same as **d** for increasing number of trials, and fitted curve 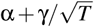 where *T* is the number of trials.

The error of the linear decoder was 1.03 ± 0.04°(s.e.m., n=6 animals) compared to 2.31°for the independent decoder (Figure 3b,c). One hypothesis for the improvements of the linear decoder is that it accounts for fluctuations in multiplicative gain, which make up a large fraction of the trial-to-trial variability ^33;50;51^. This is unlikely to be true, because adding a trial-by-trial multiplicative gain to the independent-neuron model only improved its performance very little (2.20 ± 0.13°, s.e.m., n=6, Figure S2a-c). The linear decoder also outperformed a discriminative generalization of the population vector decoding approach ^35;52^ (2.77 ± 0.15°, s.e.m., n=6, Figure S2d-f).

The best decoding was achieved for horizontal gratings, and the worst for vertical gratings, although the differences were small (Figure S3). The linear decoder was not affected by the presence of spontaneous activity patterns because it performed equally well whether these patterns were subtracted out or not (Figure S4). This confirms our previous results, that stimulus responses live in an orthogonal space to the spontaneous, behaviorally-driven activity ^33^. Finally, the lowest decoding error could not be achieved after reducing the dimensionality of the data to fewer than 128 principal components (Figure S2g-i).

To determine if the linear decoder achieved the minimum possible error, we analyzed the asymptotic behavior of the decoding error. We found that the error continued to decrease with increasing numbers of neurons and stimuli, and estimated that it would plateau at 0.71 °. in the limit of very many neurons (Figure 3d), and 0.38 °. in the limit of very many training set stimuli (Figure 3e). This analysis suggested that the most effective way to further improve decoding performance would be to show more stimuli, presumably because this would further prevent overfitting.

### Neural discrimination thresholds of 0.3°

We therefore ran such experiments next, by restricting the range of stimuli to 43-47°, and presenting ∼ 4,000 stimuli in this range. This increased the stimulus density from ∼ 10 trials/deg to ∼ 1,000 trials/deg. To avoid boundary effects when decoding orientations from such a limited range, we switched to a decoding task in which the decoder reports if the true stimulus is above or below the middle value of 45°. For this new decoding task, we fit linear decoders to directly predict the difference in angle between the presented stimulus and the middle stimulus of 45°, and evaluated on test data if the decoder correctly reported whether the stimulus was larger or smaller than 45 °(Figure 4a).

**Figure 4:**
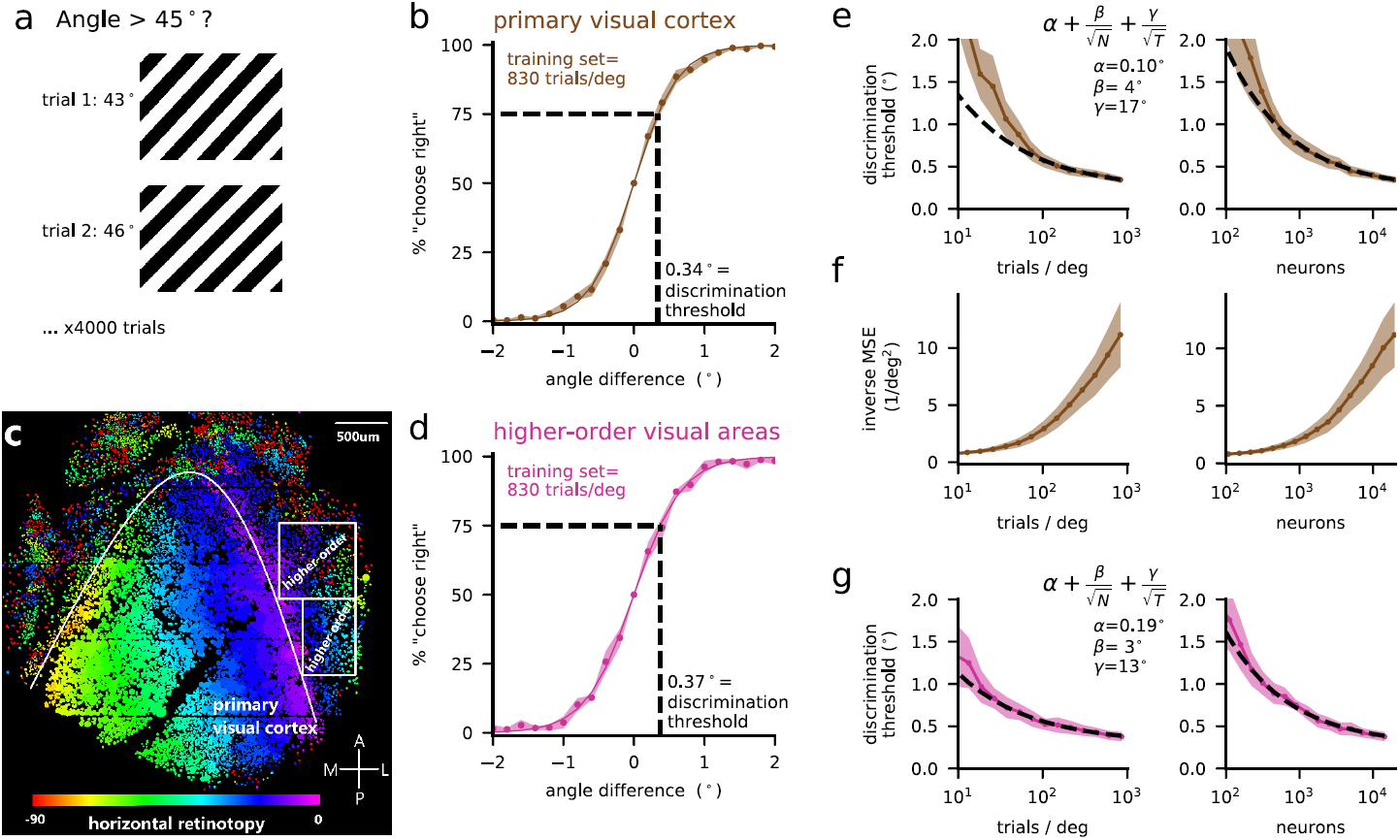
Orientation discrimination in primary and higher-order visual cortex. **a**, Decoding task schematic. Stimuli in the range 43-47° were passively shown to the mice, and the neural activity was used to decode if the stimulus angle was smaller or larger than 45°. **b**, Neurometric curves were symmetrized (see Methods) and averaged across sessions. A sigmoid function was fit to the neurometric data points. The discrimination threshold was computed from the sigmoid as the angle difference for which 75% correct performance is achieved. **c**, Horizontal retinotopy for single cells recorded with a mesoscope. The lateral higher-order visual areas with reversed retinotopy were targeted for multi-plane recordings. **d**, Same as b for higher-order areas. **e**, Discrimination thresholds as a function of the number of trials and neurons used for decoding, averaged across sessions. Also plotted are fits of the function 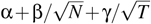 to the asymptotic behavior of the decoding error. **f**, The inverse mean-squared error (MSE), from predicting the stimulus with a linear decoder, as a function of trials / deg and neurons. **g**, Same as **e** for higher-order areas.

Following common conventions, we defined the neural discrimination threshold as the angle at which the decoder achieved 75% correct performance. We found an average discrimination threshold of 0.34 ± 0.04°(s.e.m., n=5 animals) for this stimulus set of 1,000 trials/deg (Figure 4b). As a sanity check, we performed control recordings with the laser path blocked (shutter on) and found no ability to decode from the very low bleedthrough of the screen into the microscope (Figure S5).

In contrast to the very low neural discrimination thresholds, reported *behavioral* discrimination thresholds in mice at 75% correct are in the range of 20-30 °. when directly estimated ^40^ or inferred from available data ^39;41^. Why then is neural decoding from primary visual cortex (V1) so much better? One hypothesis is that the information never leaves the primary sensory area. To test this, we repeated the experiments in higher-order visual areas that are lateral to V1, identified by retinotopic mapping at single cell resolution (Figure 4c). Again we found very low discrimination thresholds of 0.37 ± 0.05 ° (s.e.m., n=3 animals, Figure 4d).

To determine if the decoding error achieved asymptotes at these values, we computed discrimination thresholds for increasingly larger, random subsets of neurons and trials (Figure 4e). We then designed a parametric model of the error as a function of the number of trials and neurons, which provided excellent fits of the asymptotic behavior of the decoding error (Figure 4e). We used this parametric model to extrapolate the decoding error from primary visual cortex to an asymptote of 0.1°in the limit of very many neurons and trials. We also calculated the inverse mean squared error of the decoder (Figure 4f), which is proportional to a lower bound on the information content by the Cramer-Rao bound ^53^. We observed no saturation in this metric either. Finally, we repeated the fitting procedure for the data acquired in higher-order visual cortex, and again found that the likely asymptote was not yet reached (Figure 4g).

For comparison, we also performed the discrimination using the original stimulus set from Figure 2 and Figure 3, which only contained a stimulus density of 10 trials/degree. For this analysis, we picked 12 boundary stimuli uniformly in the range 0-360°, and discriminated stimuli within 30 °. The discrimination thresholds were 1.06 0.04°. (Figure 5a), somewhat lower than the 2°.estimate for the same number of trials using the high density stimulus set (Figure 4d). This discrepancy may be explained by the higher rates of response adaptation in the latter case.

**Figure 5:**
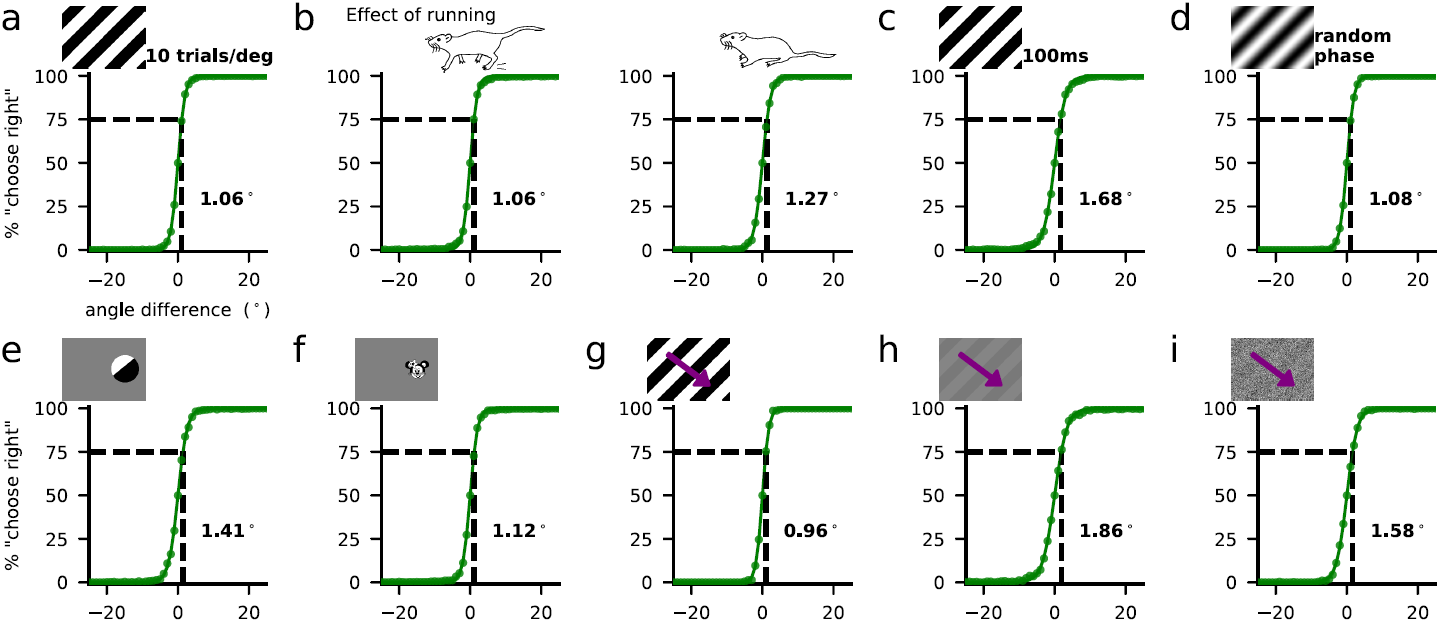
Discrimination thresholds for different stimuli and behavioral state changes. **a**, Neurometric curve for data from Figures 2 and 3, divided into intervals of ± 30°, and analyzed with a discrimination decoding task like that in Figure 4. Note the much smaller density of trials (10 trials/deg vs 1,000 trials/deg). The line is a sigmoid fitted to the data points. Error bars across recording sessions (s.e.m.) are plotted, but they are smaller than the marker size. **a**, Neurometric curves for subsets of trials divided into running and passive trials. **c-i**, Neurometric curves for other stimulus types: **c**, short, **d**, sinusoidal and random phase, **e**, localized, **f**, complex, **g**, drifting, **h**, drifting + low-contrast, **i**, drifting + low-contrast + noise. All these stimulus sets were presented at ∼10 trials/deg.

We wondered if some of the difference between neural and behavioral discrimination could be related to behavioral states, because the mice in our experiments were free to run on an air-floating ball. To quantify running-related differences, we split test trials (but not training trials) in two groups based on locomotion, and found that passive trials had modestly increased discrimination thresholds of 1.27°. compared to 1.06°. for running trials (Figure 5b). Thus, behavioral states cannot account for the large discrepancy between behavioral and neural discrimination thresholds.

We next asked if the discrepancy might be accounted for by stimulus properties. We recorded neural responses to new stimulus sets (Figure 5c-i) to investigate this possibility. We varied the duration of the static grating stimuli from 750ms to 100ms (Figure 5c), and changed it from a square to a sinusoidal grating, varying the phase randomly on every trial (Figure 5d). We also varied the size of the static grating stimuli from full field to 30°. (Figure 5e), and we presented a complex stimulus without a well-defined absolute orientation that was rotated around its center (Figure 5f). Finally, we presented drifting gratings (2Hz), drifting gratings with low contrast (5%) and drifting gratings with low contrast and large added noise (Figure 5g-i). These manipulations either did not increase discrimination thresholds or did so modestly, up to at most 1.86°.for the low-contrast, drifting gratings (Figure 5h).

We also considered the possibility that the neural code might change over time, making decoders obtained from the first half of the recording ineffective on the second half. We therefore split train/test trials chronologically rather than choosing them interspersed throughout the recording. We found a modest increase in discrimination threshold to 1.17°. Compared to the original 1.06°. (Figure S6a,b). We also did not find a difference in discrimination threshold between layers 2/3 and layer 4 (Figure S6c,d).

Thus, neither the stimulus properties nor the behavioral states can account for the discrepancy between behavioral and neural discrimination thresholds. We conclude that mice are not using the available neural information efficiently in laboratory tasks. There may be several reasons for this, which we explore in the discussion. First, we show that the discrepancy between neural and behavioral discrimination is not necessarily a consequence of trial-by-trial learning limitations.

### Biologically-plausible learning by perceptrons

It might be that the sequential nature of trial-by-trial learning makes it difficult to learn optimal decoders without storing the neural responses to all previously presented stimuli, which seems unfeasible. We constructed online decoders which process trials sequentially by using perceptrons ^53;54;55;56;57^ (Figure 6a). The perceptron sums the activities of its input neurons **x** multiplied by a set of weights **w** (*y*_pred_ = **w · x**). The sign of *y*_pred_ is then used to predict the label (−1,+1).

**Figure 6:**
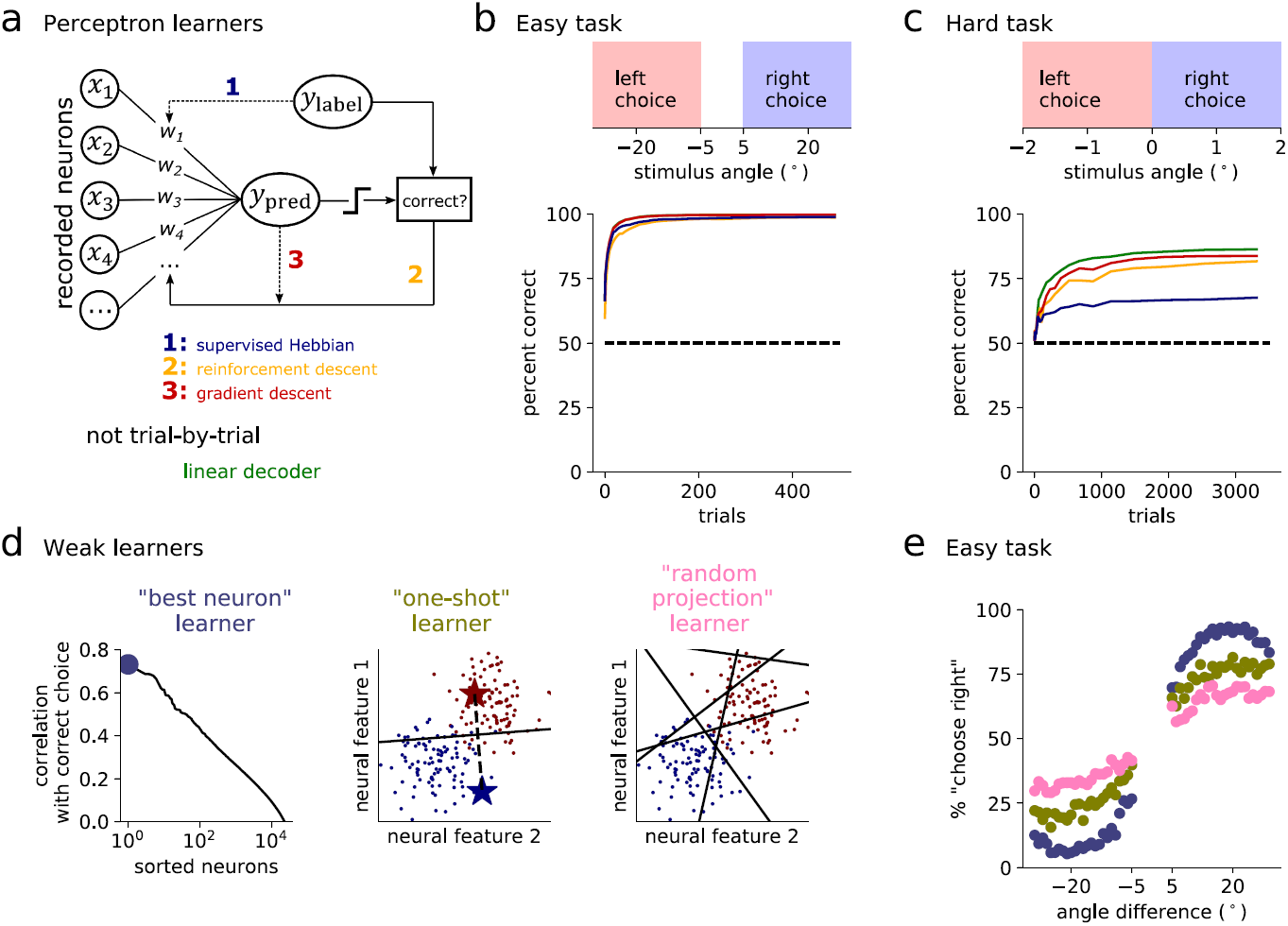
Online learning of discrimination tasks. **a**, Trial-by-trial learning for a perceptron can use either 1) only the label information (blue), 2) only the trial reinforcement (yellow), or 3) both the trial reinforcement and the continuous valued prediction (red). Finally, non-incremental learning can use all trials simultaneously, such the linear decoder previously used in Figure 3 (green). **b**, “Easy task” configuration and decoder learning performance, averaged over the neural datasets which provide the inputs **x. c**, Same as **b** for the “hard task”. **d**, Alternative learning strategies that are plausible but weak. The “best neuron” learner chooses the neuron most correlated with the correct choice on the training set. The “one shot” learner uses just one training example of each class (left and right) to find a linear discriminant. The “random projection” learner chooses the best performing random projection out of 100. **e**, Performance of the weak learners on the easy task.

The objective of learning in a perceptron is to change the weights so as to minimize the mean-squared error of the prediction.

Simple forms of online learning in a perceptron can be biologically realistic if they only require global error signals in addition to information available locally at a synapse. We investigate three such versions here. First, we consider a “supervised Hebbian” learner, which changes the weight of neuron *k* (*w*_*k*_) using the trial label *y*_label_, which is either −1 or +1, and the neural response *x*_*k*_ (Figure 6a):

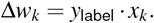

We also consider a gradient descent learner, which changes the weights by

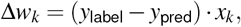

and a restricted form of gradient descent called “reinforcement descent”, which uses the reinforcement feedback (correct/wrong) in place of the full prediction error:

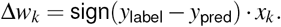

To test these online learning strategies, we designed two tasks, one easy and one hard. In the easy task, the learning agents used the neural responses to our first stimulus set with 10 trials/deg, restricted to angles between −30 and −5 degrees for the “left” choice, and 5 to 30 for the “right” choice (Figure 6b). All three perceptron learners performed the easy task perfectly, using a small number of training trials (Figure 6b). In the hard task, the learners had to discriminate between positive and negative stimulus angles of up to 2 °., using the neural data recorded at 1,000 trials/deg (Figure 6c). The supervised Hebbian learner was unable to perform well in this task with an asymptotic performance of 66%. However, the learners based on gradient descent and reinforcement descent performed relatively well at 83% and 81% compared to 85% for the optimal linear decoder, which had instantaneous access to all trials in its history (Figure 6c). Therefore, the perceptrons that used error feedback acquired information nearly as efficiently as the optimal linear decoder did.

We conclude that simple online decoders can learn even difficult orientation discrimination tasks from neural data in a sample-efficient way. Therefore, sequential learning strategies do not fundamentally limit behavioral task performance. It follows that animals use suboptimal learning strategies in laboratory experiments, perhaps because those strategies are favorable in ecological contexts. We end this study by proposing examples of weak but potentially relevant decoders, and leave it to future work to evaluate these weak learners against animal behavior. The decoders we propose (Figure 6e) are: 1) the “best neuron” learner, which finds the neuron most correlated with the correct choice on the training set and uses this neuron to make predictions on the test set; 2) the “one-shot” learner, which learns from only one trial of each stimulus category and builds a linear decoder along the discriminant direction of these two trials; and 3) the “random projection” learner, which evaluates 100 different random projections and retains the one that maximizes training set performance. These decoders had test set performance on the easy task in the 65-90% range (Figure 6e), which is in line with mouse behavioral performance on similar tasks ^39;40;41;58;59^.

## Discussion

Here we have shown that mouse visual cortex encodes visual features to a very high precision on a single-trial basis, despite large single-neuron variability and multi-neuron co-variability. To show this we recorded from large populations of neurons in visual cortex and used decoders to compute neurometric discrimination thresholds as low as 0.34° in mice, a species thought to have poor visual acuity. We also decoded the visual stimulus from higher-order visual areas and found similarly low thresholds of 0.37°. Simple linear decoders were sufficient to achieve this performance, although it is possible that nonlinear decoders may perform even better ^60^ (but see our analyses of multi-layer networks and random forest classifiers in Figure S6e,f). Furthermore, the neural discrimination thresholds did not appear to saturate with the number of neurons and trials, leading to asymptotic estimates of ∼0.1°. The neural discrimination thresholds we estimated were ∼ 100x lower than the behavioral discrimination thresholds reported by previous studies ^39;40;41^. We could not explain the difference between behavioral and neural discrimination by varying several stimulus properties, by splitting trials according to behavioral states, or by learning the decoders in a biologically plausible way. Our results imply that neural noise does not set a fundamental limit on the accuracy of sensory perception.

One previous study in primates has found that the neurometric detection sensitivity exceeded the animal’s behavior ^61^, however by a much smaller amount than what we have shown here (less than 2x contrast threshold improvement). Furthermore, the 0.3 degree neural discrimination thresholds we report in mice are lower than even the reported thresholds of highly-trained humans in this task, which were estimated at ∼1° ^62^. We suggest three potential explanations for the discrepancy between behavioral performance and neural information content. First, it is possible that downstream decoders in the mouse brain are computationally limited, and cannot find the same discrimination directions in neural space that are found by an ideal external observer ^63^ (Figure 5). Second, it is possible that the fine stimulus information is not available for perceptual report, but is used by other visual functions such as change detection or sensory-guided motor behavior ^59;64;65^. Finally, it may be that animals do have latent task knowledge even when their behavior is poor ^66^, in which case future task improvements could target the expression of this latent knowledge, for example via immersive virtual reality navigation ^41;58;67^.

To make progress on neural coding, future computational and experimental work could focus more on ethological tasks and learning strategies ^68;69^. Learning strategies that are optimal in complex, dynamic environments may perform poorly in laboratory tasks where a very small number of stimuli are shown thousands of times to the same animal. Such tasks have been rationally designed to isolate the phenomenon of interest — sensory perception — but may have stripped away the relevant behavioral contexts which make sensory perception difficult. Perception in the real world is only one part of an animal’s strategy, and requires coordination with other neural functions like decision-making, memory and motor control. Therefore, the true neural limitations of sensory perception may only be revealed in the context of more complex and more naturalistic tasks.

## Acknowledgments

We thank Dan Flickinger for assistance with the twophoton microscope, Salvatore DiLisio and Sarah Lindo for surgeries, James Fitzgerald and Ruben Moreno-Bote for useful discussions. This research was funded by the Howard Hughes Medical Institute.

## Data and code availability

All of the processed neural activity is available on figshare ^44^(https://figshare.com/articles/Recordings_of_20_000_neurons_from_V1_in_response_to_oriented_stimuli/8279387). The code is available on github (https://github.com/MouseLand/stringer-et-al-2019a).

## Methods

### Animals

All experimental procedures were conducted according to IACUC.

We performed 18 recordings in five mice bred to express GCaMP6s in excitatory neurons: TetO-GCaMP6s x Emx1-IRES-Cre mice (available as RRID:IMSR JAX:024742 and RRID:IMSR JAX:005628). We also performed 17 recordings in three wild-type C57 mice. In these mice, AAV-GCaMP6s-P2A-nls-dTomato (RRID:Addgene 51084) was expressed virally through injections into visual cortex. These mice were male and female, and ranged from 2 to 8 months of age. Mice were housed in reverse light cycle.

### Surgical procedures

Surgical methods were similar to those described elsewhere ^33^. Briefly, surgeries were performed in adult mice (P35–P125) under isoflurane anesthesia (5% for induction, 1-2% during the surgery) in a stereotaxic frame. Before surgery, buprenorphine was administered as a systemic analgesic and lidocaine was administered locally at the surgery site. During the surgery we implanted a head-plate for later headfixation, and made a craniotomy of 4 mm in diameter with a cranial window implant for optical access. We targeted virus injections (50-200 nl, 1-3 × 1012 GC/ml) to monocular V1 (2.1-3.3 mm laterally and 3.5-4.0mm posteriorly from Bregma). To obtain large fields of view for imaging, we typically performed several injections at nearby locations, at multiple depths (∼500 *µ*m and ∼200 *µ*m). After surgery, buprenorphine was administered in the drinking water as a post-operative analgesic for two days.

### Data acquisition

We used a custom-built 2-photon mesoscope ^70^ to record neural activity, and ScanImage ^71^ for data acquisition. Multi-plane acquisition was controlled by a resonance mirror, with planes spaced 25 *µ*m apart in depth. 10-17 planes were acquired sequentially, scanning the entire stack repeatedly on average at 3 Hz. We synchronized stimulus presentation to the beginning of each frame for the first plane, and computed stimulus responses from the first three frames acquired after stimulus onset for each plane. We used a custom online Z-correction module (now in ScanImage), to correct for Z and XY drift online during the recording.

The mice were free to run on an air-floating ball. For all static image presentations an LED tablet screen at a 45° from the left eye (we recorded in right visual cortex). For drifting grating image presentations a custom circular screen made of LED arrays was placed around the head of the mouse ^72^. We also repeated the static grating experiments with this screen and obtained comparable decoding errors. To prevent direct contamination of the PMT from the screen, we placed gel filters in front of the screen which exclude green light, and used only the red channel of the screens.

For each mouse, recordings were made over multiple days, always returning to the same field of view. The field of view was selected on the first recording day such that large numbers of neurons could be observed, with clear calcium transients and a retinotopic location that was localized on the screen (identified by neuropil fluorescence responses to sparse noise). For three mice, we recorded in higher-order visual areas, located based on retinotopic responses to sparse noise. We were able to identify with high confidence the reversal in horizontal retinotopic preference, and used the random-access mesoscope to record from two fields of view simultaneous in higher-order visual areas that would correspond primarily to areas LM and PM in the Allen Brain Atlas ^73^.

### Visual stimuli

We showed various gratings and localized images. To present stimuli, we used PsychToolbox-3 in MATLAB ^74^. The stimuli were presented for 750 ms (unless otherwise stated), alternating with a gray-screen interstimulus interval lasting on average 650 ms. After every 150 stimuli, the screen was left blank (gray screen) for 32 seconds. The activity during these non-stimulus periods was used to project out spontaneous dimensions from the neuronal population responses (see below).

All gratings shown had a spatial frequency of 0.05 cycles / degree. Almost all stimuli were square gratings at a fixed phase, except for the sinusoidal-intensity gratings, which also had a random phase. All stimuli had full contrast, except for the low contrast and low contrast+noisy drifting gratings, which had a contrast of 5%. Drifting gratings had a temporal frequency of 2 Hz. We showed random orientations of these stimuli between 0° and 360° on each trial. For the stimuli with random phase, there is no difference between orientations that are 180° out of phase, therefore we pooled those trials, resulting in a range of orientations between 0° and 180°.

The localized stimulus was restricted to 30° of visual space and used a lower frequency grating, so that it contained only one black and one white sub-field. The complex “minimouse” stimulus also spanned 30° of visual space and was rotated around its center. Outside of these stimuli, the screen was gray.

### Calcium imaging processing

The calcium imaging processing pipeline and the subsequent analyses use numpy, scipy, numba, scikitimage, and scikit-learn ^75;76;77;78;79^. The figures in the paper were made using matplotlib in jupyter ^80;81^.

Calcium imaging data was processed using the Suite2p toolbox ^42^, available at www.github.com/MouseLand/suite2p. Suite2p performs motion correction, ROI detection, cell classification, neuropil correction, and spike deconvolution as described elsewhere ^33^. For non-negative deconvolution, we used a timescale of decay of 1.25 seconds ^43;82^. We obtained 18,496 ± 3,441 (s.d., n=35) neurons in the recordings.

In three recordings, we closed the shutter on the 2p laser. In this case the extracted ROIs were randomly placed disks with similar numbers of average pixels as measured for real cells. These traces were neither neuropil-corrected, nor deconvolved.

### Stimulus responses

We defined the stimulus response as the summed activity of the first three bins (∼1 second) following stimulus onset. We split the trials 75/25 into training and testing sets, with every 4th trial assigned to the test set.

### Splitting cells into two populations

When looking at correlated decoding errors, we split the neurons into two populations. We first divided the XY plane into 8 non-overlapping strips of width 150 *µ*m, and assigned the neurons in the even strips to one group, and the neurons in the odd strips to the other group, regardless of the neuron’s depth. Thus, there did not exist neuron pairs in the two sets that had the same XY position but a different depth. This special division was performed to avoid contamination artifacts between overlapping cells or between consecutive planes.

### Tuning curves and SNR

We fit the training trials with *n*_basis_ = 10 cosine and sine basis functions, where θ is the angle of the stimulus shown in radians:

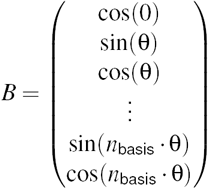

We performed linear regression from *B* to the neural responses and used the fitted function *f*_*n*_(θ) as the tuning curve. To compute the signal-to-noise ratio, we defined the signal as the variance of the tuning curve and the noise as the variance of the residual noise after subtracting out the tuning curve value.

### Removal of ongoing activity dimensions

Approximately a third of the variance of visual cortical population activity represents behavior-related fluctuations ^33^. In Figure S4 we projected out the ongoing activity dimensions to show that they had no influence on sensory coding. To do this we computed the top 32 principal components of z-scored and binned (3 frames ≈ 1 second) ongoing activity. The spontaneous, ongoing neural activity was recorded during gray screen presentations, and these were presented after every 150 stimulus presentations for 32 seconds each time. To remove these dimensions from stimulus responses, the stimulus-driven activity was first z-scored (using the mean and variance of each neuron computed from ongoing activity), and then we computed the projection onto the 32 top spontaneous dimensions and subtracted it off from the z-scored stimulus-driven activity (Figure S4, y-axis).

### Independent decoder

For the independent decoder, we built generative models of the neural data by modelling the mean and standard deviation of each neuron on the training set. The mean was obtained as a function of stimulus angle using the tuning curve fits *f*_*n*_(θ) above. The standard deviation σ_*n*_(θ) was fit similarly, after subtracting the mean predictions on the training set, by squaring the residuals and fitting them in the same set of basis functions. With the mean and standard deviation defined for each neuron and each stimulus, we computed the probability that a novel neural pattern *R*(*n*) was produced by a putative stimulus orientation θ′:

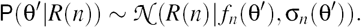

These probabilities were evaluated in log-space for a discrete set of θ′ (n=48 orientations) and summed across neurons. To decode orientations more finely, we upsampled the log-probability curves by a factor of 100 using kriging interpolation. The stimulus angle corresponding to the peak of the upsampled curve was used as the decoded orientation.

We also tested an extension of this decoder with multiplicative single-trial gains. On single trials, we determined the best-fitting multiplicative gain *g*_*n*_ and used it to compute the likelihood:

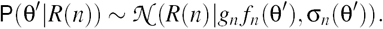

### Linear decoder

To decode circular angles using locally linear decoders, we regressed the neural activity onto “super”-neurons (see Figure 3a). These super-neurons were von Mises tuning curves (*n* = 48) with peaks equally spaced along 360° and with σ = 0.1:

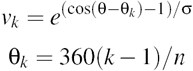

We fit the transformation from neurons to superneurons on training trials using a ridge regularization constant of 1 (the default for the scikit-learn Python package), and then predicted the super-neuron responses on test trials. We then upsampled the superneuron responses from 48 to 4,800 to decode stimulus angle more finely (like we did in the independent decoder). For efficient estimation of the linear decoder we used the matrix inversion lemma in the case where the number of trials was less than the number of neurons.

### Conditional probability model

In Figure S2 we evaluate a conditional probability model, which can account for correlations, just like the linear models shown in the main text. The model is a generalization of the population vector approach and follows in the footsteps of previous work in this direction ^35;52^. We fit the model directly by maximum likelihood estimation, unlike previous methods which used indirect fitting procedures. The conditional probability model is defined implicitly by its energy function

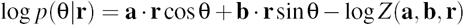

where **r** is the vector of neural activity in response to a stimulus, **a, b** are parameters to be learned from the training data, and *Z* is the partition function of this energy model, which ensures that probabilities normalize to 1. The advantage of this model is that it is defined probabilistically and has fewer parameters than the linear decoders. However, fitting **a, b** is analytically intractable. We adopt a gradient descent optimization strategy, which requires numerical integration of the log-partition function over the 1-dimensional orientation variable:

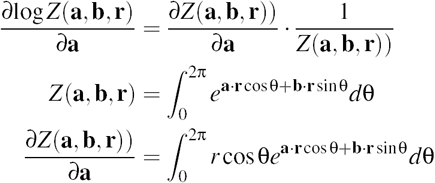

and an equivalent derivation for ∂log(*Z*)*/*∂*b*. Thus, on each step of gradient descent, numerical integrals must be computed for *Z*, ∂*Z/*∂*a* and ∂*Z/*∂*b*, for each sample **r** of neural activity on the training set. The gradients for the rest of the log *p*(θ| **r**) are straightforward. On test data, the most likely presented stimulus can be easily inferred as the angle of the complex number **a** · **r** + *i***b** · **r**.

We used a momentum of 0.95, and ran the optimization for 800 iterations. No further improvements on the validation set could be observed by running the optimization longer. We also tested L2 regularized versions of this model, which brought small improvements and constitute the results shown in Figure S2. The optimization code is included in the repository for this paper.

### Neurometric fitting

For the discrimination task, we used a simple linear decoder for the high density stimulus set, because the range of predicted angles is small (4°) and thus the behavior of the neural data is expected to be linear in that range. For the low density stimulus set, we predicted angles in the range ± 30° by first predicting a function *f* of the stimulus θ and the threshold stimulus θ_0_, which was a difference of von Mises basis functions:

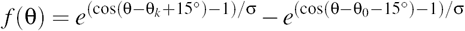

where σ was the same as it was for the linear decoder. We used the sign of the neural prediction *f* on test trials as the predicted class label of the decoder.

To quantify discrimination performance, we computed the probability *P*(θ) as the fraction of correct predictions when θ was in a bin of 1° (10 trials/deg stimuli) or a bin of 0.1° (1,000 trials/deg stimuli). This produced the neurometric data points. For the 10 trials/deg stimuli, we used 32 different threshold stimuli (θ_0_), spaced evenly from 0° to 360°, and averaged the neurometric curves from these 32 different discrimination tasks. To symmetrize the neurometric curves, we used

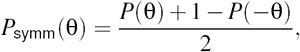

as if the discrimination task was repeated with left vs right labels interchanged. To fit *P*(θ) we used centered sigmoids with a single free parameter β:

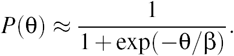

We fit β using the “curve fit” function in scipy ^76^. The fit curves are shown as continuous lines in Figure 4b and Figure 5. To compute the discrimination threshold, we took the value of this function at 75%:

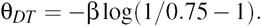

### Asymptotics

To fit the asymptotic error, we modeled the scaling of the median error with the parametrization 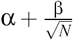, where *N* is the number of neurons or 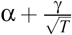, where *T* is the number of trials (Figure 3d,e). The scaling of 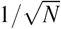 was chosen because it corresponds to the decay of the standard deviation of an average of independent random variables with the same variance. We fit α and β (or α and γ) to the last 12 points of each curve in Figure 3d,e using linear regression.

In Figure 4e we fit the function 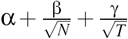, where *N* is the number of neurons and *T* is the number of trials per degree (4 degrees total shown in these experiments). We fit this function using linear regression to the discrimination thresholds obtained with data subsampled at *>* 100 trials/deg and *>* 1, 000 neurons (72 points in total).

### Neural networks for decoding

We used PyTorch ^83^ to train a neural network to perform the discrimination task in Figure 3 on the 10 trials/degree recordings. This network consisted of two rectified-linear layers and an output sigmoid. The input layer consisted of 256 units using the top principal components from the data (this reduced overfitting). The two hidden layers consisted of 100 units each. We trained the network for 50,000 iterations using stochastic gradient descent with momentum of 0.9 and a learning rate of 1e-3. The cost was a binary cross-entropy loss. We averaged over 5 random initializations of the neural network for each recording. Figure S6e shows the average discrimination performance over all recordings.

### Random forests for decoding

We used scikit-learn ^79^ to train a random forest ensemble classifier to perform the discrimination task using the neural activity from the 10 trials/degree recordings. We used 1000 trees and averaged over 5 random initializations of the classifier for each recording. Fig-ure S6f shows the average discrimination performance over all recordings.

### Perceptron learners

The perceptrons used the learning rule for weight *w*_*k*_ for neuron *k*:

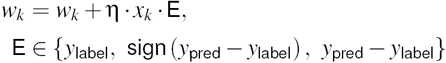

where η is the learning rate and *y*_pred_ = **w** · **x**. The three cases for E correspond to the supervised Hebbian, the reinforcement descent and the gradient descent decoders. The gradient descent decoder has been derived from the error cost function:

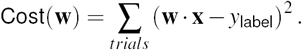

### Signal variance

We computed the signal variance of the static gratings responses using two repeats of the same stimuli ^38;43^. We estimated the signal variance as the correlation coefficient between the two repeats - the “repeat correlation”. Because the stimuli we presented were of random orientation, we technically did not have two repeats of the exact same stimulus. Therefore we sorted the stimuli by orientation and divided the responses into two halves using this sorting: one half was the odd stimuli and the other half was the even stimuli. The average repeat correlations over all neurons in 6 recordings was 0.11.

### Analysis of electrophysiological data

We quantified the SNR of mouse electrophysiological recordings collected by the Allen Institute using Neuropixels probes ^46^. We used the recordings in which the “brain observatory 1.1” stimulus set was shown. During these recordings a variety of stimuli were shown including static gratings with 4 possible phases, 6 possible orientations, and 5 different spatial frequencies presented for 400 ms each. We used the responses of neurons to the static gratings at spatial frequency 0.04 cpd (because this was closest to our gratings of 0.05 cpd). The response of each neuron was defined as the sum of spikes in a 400 ms window following the stimulus onset. We then divided the responses into two halves, matching the phases in each half, resulting in two sets of at least 123 responses each (it varied slightly from dataset to dataset). The repeat correlation of each neuron was computed in the same way as it was for the two-photon calcium imaging data, using the correlation coefficient between the two repeats. The average repeat correlations over primary visual cortex neurons in 32 recordings was 0.12.

---

**S1:**
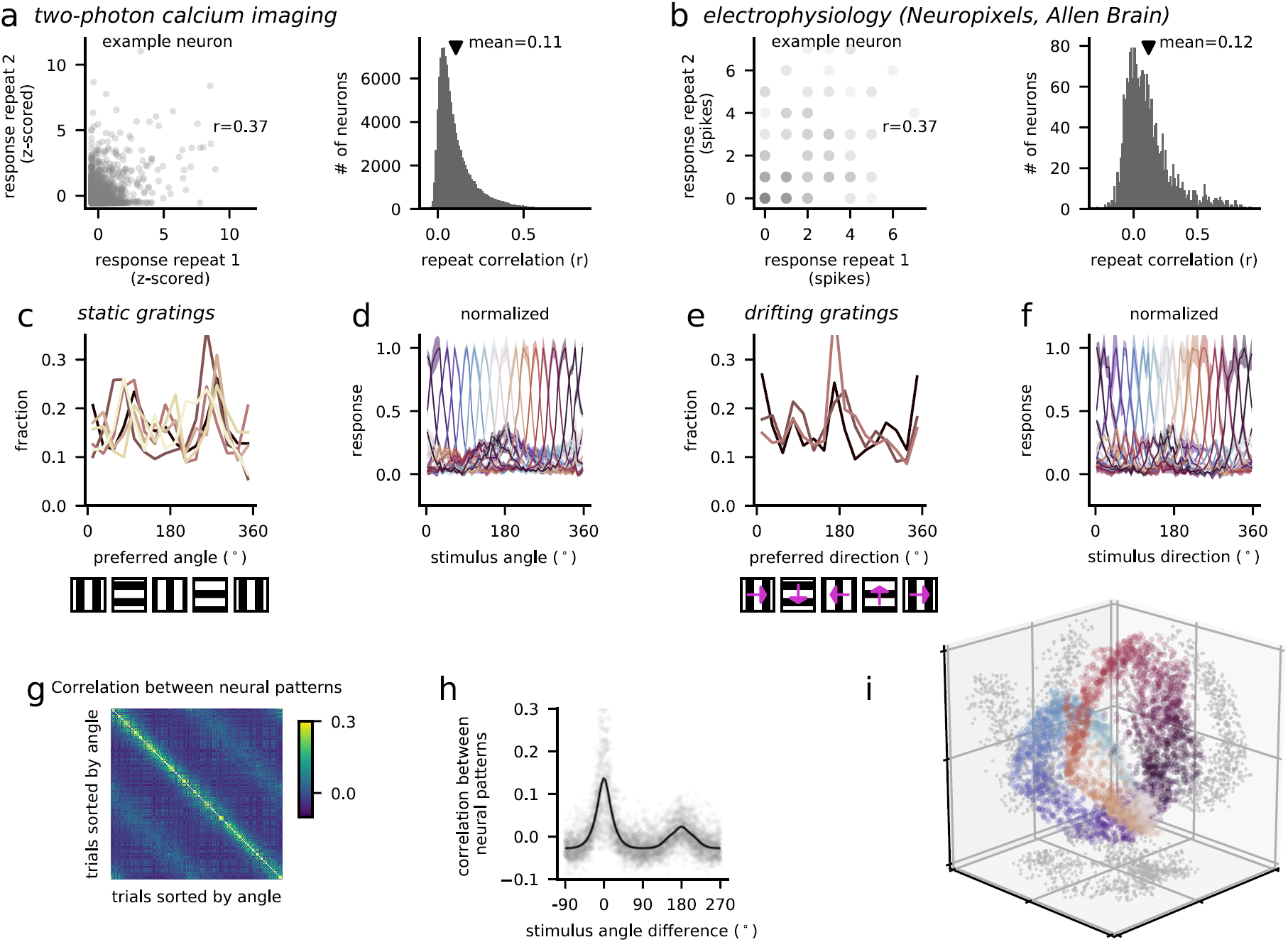
Signal variance and tuning curves. **a**, Signal variance from two-photon calcium imaging of responses to static gratings (0.05 cpd) at fixed phase (0°). Left: example neuron responses to two repeats of the same orientations and correlation *r* between responses to the two repeats. Right: Distribution of *r* across neurons in our recordings. *r* is also an unbiased estimate of the signal variance in each neuron’s responses. **b**, Same as **a** for electrophysiological recordings using Neuropixels in primary visual cortex from 46. Static gratings of 6 different orientations at a similar spatial frequency (0.04 cpd) and 4 different phases were shown for 400 ms each. The response is the sum of spikes in the 400 ms following the stimulus onset. **c**, Distribution of preferred orientation across cells. Each line represents a different recording. **d**, On test trials, the responses of neurons with similar preferred angles (determined on training trials) were averaged across all recordings, and normalized between 0 and 1. The half-width half-max of these tuning curves was 14.1°. **e**,**f**, Same as **c**,**d** for drifting grating responses. The half-width half-max of the tuning curves in **f** was 15.1°. **g**, Correlation between neural responses on all pairs of trials of static grating recordings. **h**, Same data as **g**, with correlation plotted as a function of stimulus angle difference. The black line shows the binned and smoothed average. **i**, The ∼20,000-dimensional response vectors were embedded into three dimensions using ISOMAP. Points are colored by the angle of the presented stimulus.

**S2:**
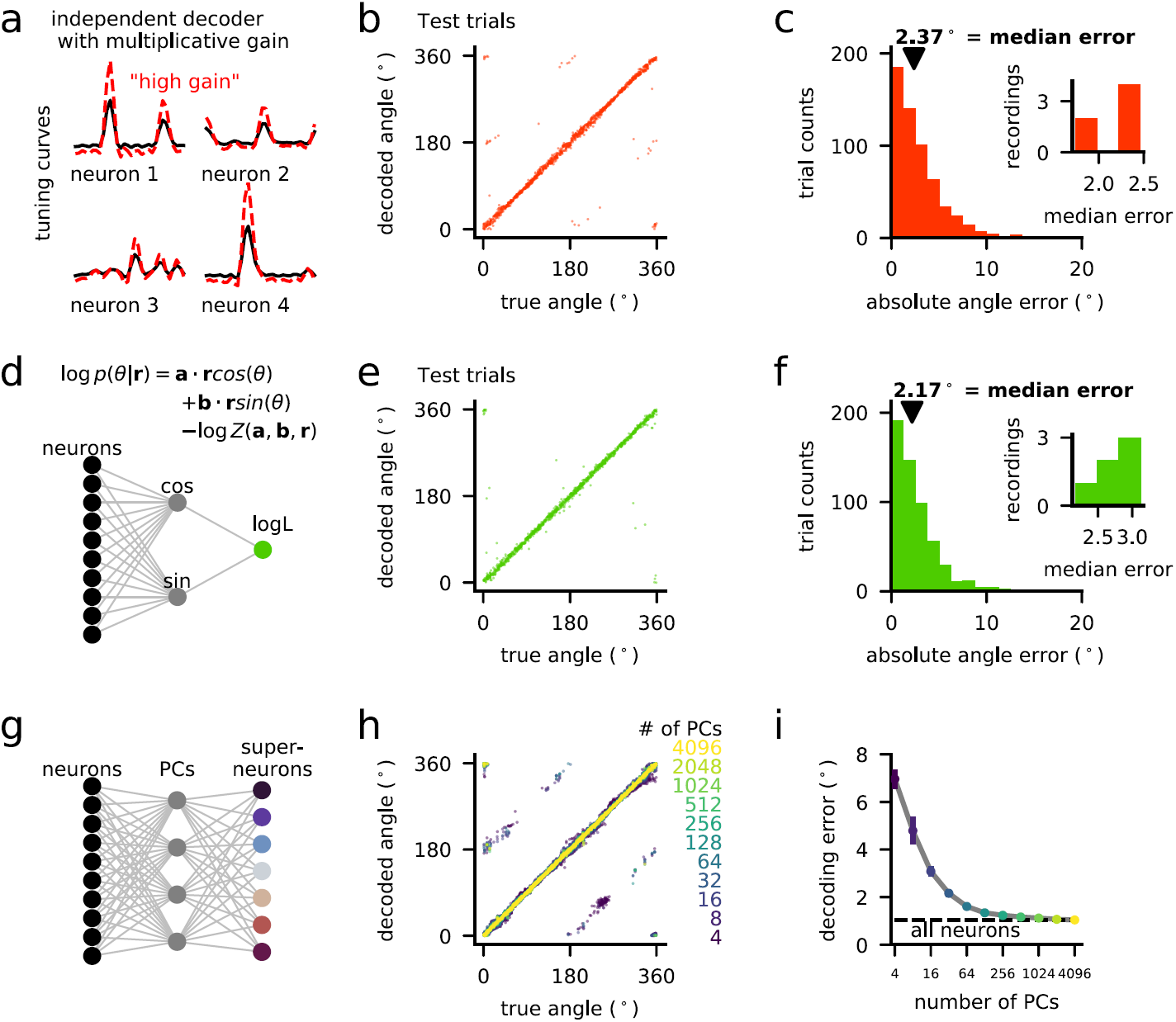
Conditional probability model, and linear decoding after dimensionality reduction. **a**, Schematic of independent decoder with multiplicative single-trial gain shared across all neurons. **b**, Decoded stimuli. **c**, Decoding errors for this model. Compare to Figure2c. **d**, Schematic of conditional probability model of the predictive likelihood distribution. **e**, Decoded stimuli. **f**, Distribution of decoding error and (inset) distribution of median decoding errors across datasets. Compare to Figure3c. **g**, Schematic of linear decoder applied after dimensionality reduction. **h**, True angle versus decoded angle for an example recording, for different number of principal components (PCs). **i**, Median decoding error as a function of the number of PCs, averaged over recordings. Error bars are standard error. Dashed line is decoding error using all neurons (without dimensionality reduction).

**S3:**
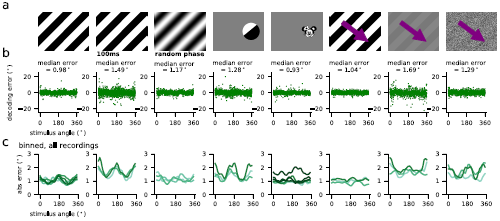
Decoding error as a function of stimulus angle. **a**, All the stimulus sets. **b**, Decoding error vs stimulus angle for each trial of an example recording of each stimulus set. **c**, Plot of B after taking the absolute value and binning. Each line is a different recording.

**S4:**
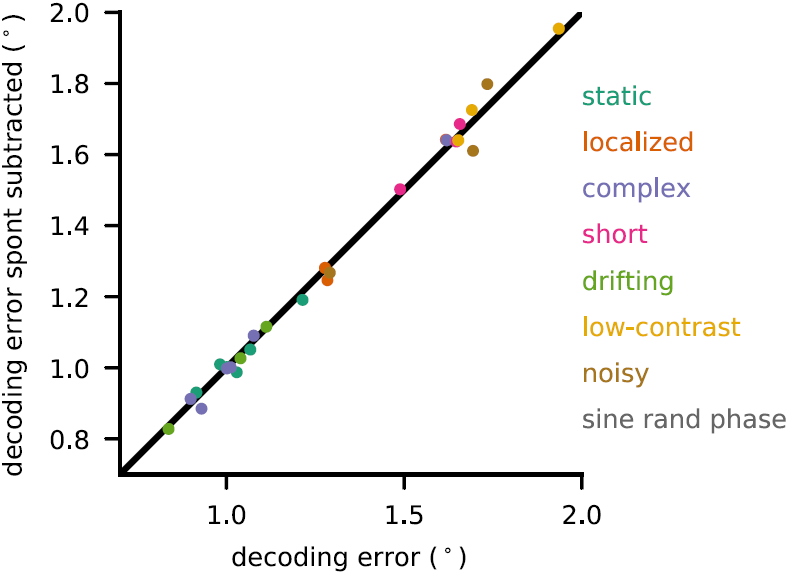
Principal components of spontaneous, ongoing activity do not influence decoding accuracy. 32 principal components of spontaneous, ongoing activity were subtracted from the stimulus responses, and the linear decoder was trained on these responses. The median decoding errors of the subtracted responses are plotted vs the original.

**S5:**
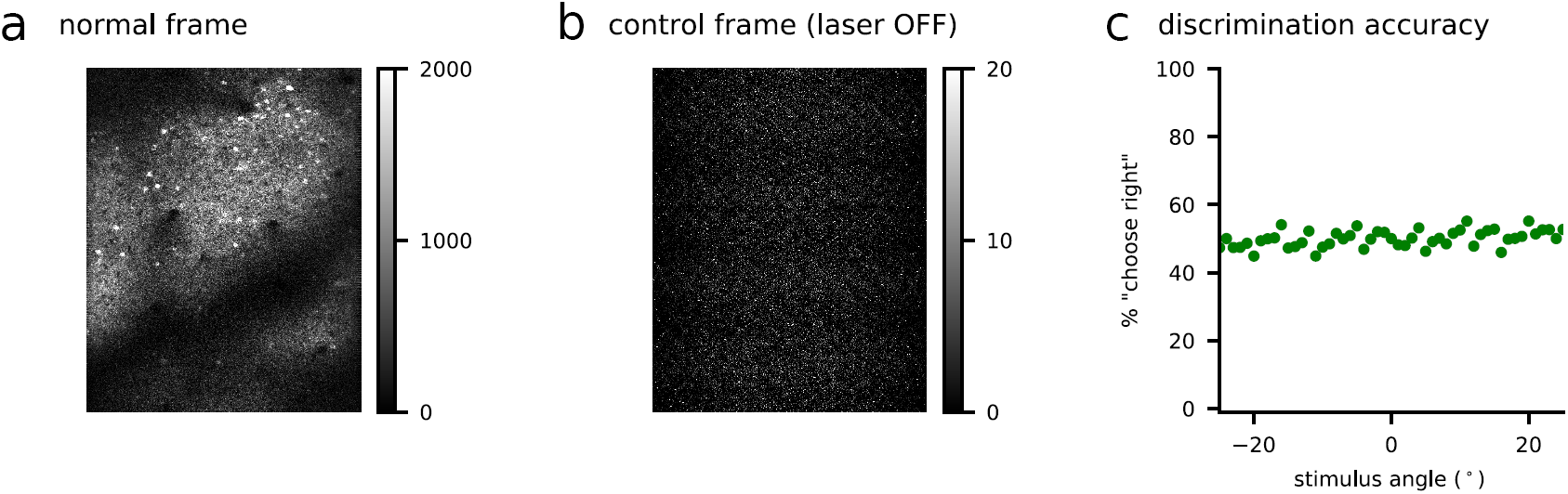
Control (laser off). To ensure that the visual stimulus screen did not contaminate the fluorescence signals collected by the photo-multiplier tube (PMT), we performed recordings in which the shutter on the 2p laser was kept closed, but all other conditions remained the same. With the laser off, the PMT signals were near-zero and reflected mainly auto-fluorescence, photon noise and 60Hz signals. **a**, Example frame from one of the recordings used in the paper. **b**, Example frame of control recording with laser OFF. **c**, Discrimination of stimuli in laser OFF recordings (n=3 mice, *>*4000 total trials/mouse). The discrimination performance appeared to be at chance.

**S6:**
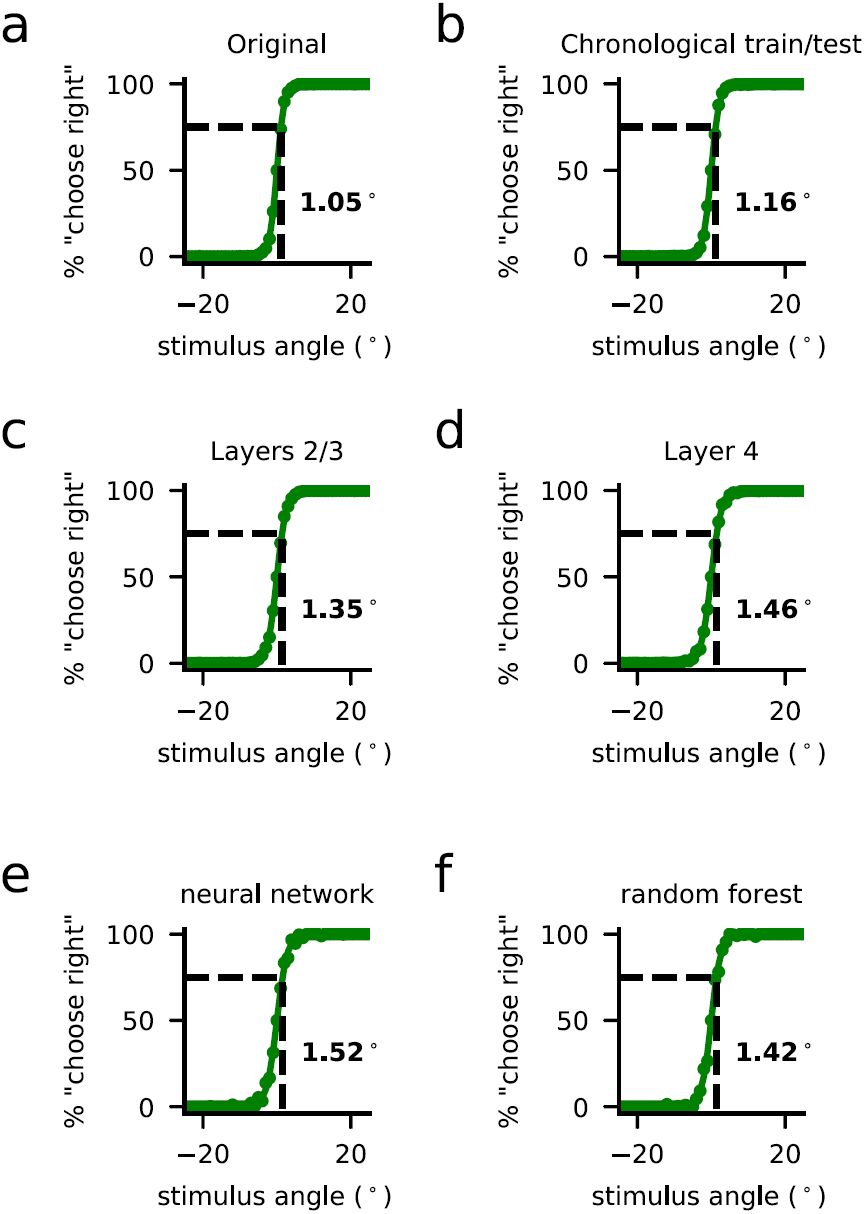
Discrimination thresholds for different data splits and decoders. **a**, Train and test trials interleaved (same data as Figure5a). **b**, Chronological split across the 120-180 minutes of recording: training trials were first 75% of stimuli presented and test trials were last 25% of stimuli presented. Compare to **a. c**, Discrimination of static gratings (10 trials/deg) using only neurons at depths ∼125-225 *µ*m. **d**, Same as **c**, with neurons at depths ∼375-475 *µ*m. **e**, Two-layer neural network on the same 10 trials/deg data. Compare to **a. f**, Random forest classifier for the same data. Compare to **a**.

